# Molecular characterization of triple negative breast cancer formaldehyde-fixed paraffin-embedded samples by data-independent acquisition proteomics

**DOI:** 10.1101/2020.09.21.306654

**Authors:** Silvia García-Adrián, Lucía Trilla-Fuertes, Angelo Gámez-Pozo, Cristina Chiva, Rocío López-Vacas, Elena López-Camacho, Guillermo Prado-Vázquez, Andrea Zapater-Moros, María I. Lumbreras-Herrera, David Hardisson, Laura Yébenes, Pilar Zamora, Eduard Sabidó, Juan Ángel Fresno Vara, Enrique Espinosa

## Abstract

Triple negative breast cancer (TNBC) accounts for 15-20% of all breast carcinomas and it is clinically characterized by an aggressive phenotype and bad prognosis. TNBC does not benefit from any targeted therapy, so further characterization is needed to define subgroups with potential therapeutic value. In this work, the proteomes of one hundred twenty-five formalin-fixed paraffin-embedded samples from patients diagnosed with triple negative breast cancer were analyzed by mass spectrometry using data-independent acquisition. Hierarchical clustering, probabilistic graphical models and Significance Analysis of Microarrays were used to characterize molecular groups. Additionally, a predictive signature related with relapse was defined. Two molecular groups with differences in several biological processes as glycolysis, translation and immune response, were defined in this cohort, and a prognostic signature based on the abundance of proteins RBM3 and NIPSNAP1 was defined. This predictor split the population into low-risk and high-risk groups. The differential processes identified between the two molecular groups may serve to design new therapeutic strategies in the future and the prognostic signature could be useful to identify a population at high-risk of relapse that could be directed to clinical trials.

## Introduction

Triple negative breast cancer (TNBC) is defined by lack of expression of estrogen and progesterone receptors, as well as human epidermal growth factor receptor 2 (HER2). TNBC accounts for 15-20% of all invasive breast carcinomas. TNBC usually exhibits an aggressive behavior and is associated with high relapse and mortality rates, most relapses occurring within the first three years after diagnosis (1–4). Relapses usually affect visceral sites, such as the lung and central nervous system (CNS) (5–9). The standard treatment consists of surgery and adjuvant or neoadjuvant chemotherapy based on a combination of anthracyclines and taxanes.

TNBC is a heterogeneous disease. Genomic studies have determined the existence of molecular subtypes, (10–13) which have different clinical evolution and response to chemotherapy (14). A molecular classification of TNBC is not currently being used in clinical practice because it does not help clinicians in making treatment decisions. New and more informative prognostic biomarkers would be useful to select the most effective treatment.

In a recent publication, we have identified a proteomics-based biomarker combination to better stratify TNBC patients according to the benefits of the adjuvant chemotherapy (15). We used an initial discovery phase with a small cohort of patients (n=26), followed by a validation phase of a subset of proteins using targeted proteomics in a large patient cohort (n=114). In that study we defined a protein-based signature that predicted response to adjuvant chemotherapy. The protein signature consisted of the combination of proteins RAC2, RAB6A, BIEA and IPYR, and its classification performance was confirmed in publicly available transcriptomics datasets from independent cohorts.

Proteomics analyses based on data-independent acquisition (DIA) methods have shown an increased reproducibility of peptide quantification across multiple samples and have become the method of choice to quantify entire proteomes in large patient cohorts (16, 17). DIA methods rely on the use of several broadband isolation windows to select and fragment all detectable peptides within a sample (17) and several isolation schemes have been described to maximize specificity, sensitivity and speed (18–20). These methods have demonstrated a high quantitative accuracy and reproducibility on previous studies and they have been successfully used to characterize breast cancer subtypes using fresh-frozen tissue samples (21, 22).

In this study we relied on the high sensitivity and improved properties of data-independent acquisition methods (20) to analyze full proteomes in a large cohort of formalin-fixed paraffin embedded TNBC samples, to define molecular subgroups and stratify patients with TNBC into a group with high or low risk of relapse that improve current and future therapeutic strategies.

## Experimental procedures

### Experimental design and statistical rationale

One hundred and forty-two formalin-fixed paraffin-embedded samples from patients diagnosed of triple negative breast cancer were analyzed using a DIA+ approach (20). Neither technical replicate analyses nor control normal tissue samples were necessary due to the large size of the clinical cohort, the nature of the samples and the objectives of the study. In addition, this study was focused in the molecular characterization of the disease and its evolution instead of the carcinogenesis mechanisms (in which comparing normal and tumor tissues are necessary); therefore, normal tissue was not used as a control.

### Patient samples and clinical-pathological variables

A cohort of TNBC patients analyzed in a previous study was now analyzed by DIA+ proteomics (15, 20). A total of 136 patients diagnosed of TNBC between 1997 and 2004, and treated in daily clinical practice at two Spanish institutions (Hospital Universitario La Paz and Hospital Universitario 12 de Octubre) were identified and retrospectively analyzed. The inclusion criteria were: patients diagnosed with triple negative invasive breast carcinoma, nonmetastatic (stage I-III) disease at diagnosis, and a minimum follow-up of two years. Collected clinical variables were: TNM (based on TNM 7^th^ edition), histological grade, age at diagnosis, chemotherapy treatment, relapse, disease-free survival (DFS), and location of the first relapse. DFS was defined as the time from surgery of the primary tumor to local and/or distant tumor relapse. This project was approved by the Ethical Committees of Hospital Universitario La Paz and Hospital 12 de Octubre and all patients signed the corresponding informed consent.

### Sample processing and protein isolation

Formalin-fixed paraffin-embedded (FFPE) samples were reviewed by an experienced pathologist and only samples with at least 50% of tumor cells were selected for the study.

Proteins were isolated as described previously (15, 23). Briefly, FFPE sections were deparaffined in xylene and washed twice in ethanol. Protein extracts were eluted in 2% SDS and protein concentration was measured using MicroBCA Protein Assay Kit (Thermo Fisher Scientific). Peptide isolations were digested by trypsin and SDS was removed employing Detergent Removal Spin Columns (Thermo Fisher Scientific). Desalted peptides were solubilized in 0.1% formic acid and 3% acetonitrile. Isotopically labeled peptides were added to peptide mixes and used as internal standard for quantification.

### DIA+ data acquisition

Peptide mixtures derived from the FFPE samples were analyzed using a DIA+ method in a LTQ-Orbitrap Fusion Lumos mass spectrometer (Thermo Fisher Scientific, San Jose, CA, USA) coupled to an EASY-nLC 1000 (Thermo Fisher Scientific (Proxeon), Odense, Denmark). Peptides were loaded directly onto the analytical column and separated by reversed-phase chromatography using a 50-cm column with an inner diameter of 75 μm, packed with 2 μm C18 particles spectrometer (Thermo Scientific, San Jose, CA, USA).

Chromatographic gradients started at 95% buffer A and 5% buffer B with a flow rate of 300 nl/min for 5 minutes and gradually increased to 22% buffer B and 78% A in 109 min and then to 35% buffer B and 65% A in 11 min. After each analysis, the column was washed for 10 min with 10% buffer A and 90% buffer B. Buffer A: 0.1% formic acid in water. Buffer B: 0.1% formic acid in acetonitrile.

The mass spectrometer was operated in positive ionization mode with an EASY-Spray nanosource with spray voltage set at 2.4 kV and source temperature at 275 °C. The instrument was operated in data-independent acquisition mode, with a full MS scans over a mass range of m/z 400–1,350 with detection in the Orbitrap (60K resolution) and with auto gain control (AGC) set to 200,000. The isolation scheme used was the same as previously reported (20). A normalized collision energy of 28% +/-5% was used for higher-energy collisional dissociation (HCD) fragmentation. MS2 scan range was set from 350 to 1850 m/z, with an AGC Target of 5.0e4 and a maximum injection time of 60 ms. The maximum injection time was set to 20 ms per segment, making a total of a maximum injection time of 60 ms per composite window. Fragment ion spectra were acquired in the the Orbitrap mass analyzer at 30K resolution.

Digested bovine serum albumin (New England Biolabs cat # P8108S) was analyzed between each sample to avoid sample carryover and to assure stability of the instrument and QCloud (24) has been used to control instrument longitudinal performance during the project.

Acquired raw data were transformed to mzXML file format with msconvert from the ProteoWizard suite v3.0.9393. Converted mzXML were further analyzed using DIA Umpire v2.1.2 (25) with the search engine Comet v2016.01 rev.0 with trypsin specificity, one allowed missed cleavage, and oxidation of methionine as variable modification (+15.9949), and carbamidomethylation at cysteine as fixed modification (+57.0214). Error tolerance was set at 10 ppm for MS1 and 0.02 Da at MS2. The swissprot human protein database with reviewed entries and decoys was used as reference database (version April 2016). Peptides and proteins identifications were filtered at 1% FDR. Peptide quantitation was based on the sum of the six most intense fragment ions. The median of all the spiked heavy peptides was used for data normalization. Protein abundances were estimated from normalized peptide abundances using the MSstats software (v3.10.6) (26). Two samples that exhibited less than 600 identified proteins were removed from the dataset for further analyses.

### Analyses of clinical variables

A descriptive analysis of the clinical parameters was performed. Statistical comparisons were done using Chi-squared test and t-test. For survival analyses, Kaplan-Meier and log-rank methods were used. All these analyses were done in SPSS IBM Statistics v20.

### Hierarchical cluster and Significance Analysis of Microarrays

Both hierarchical cluster (HCL) and Significance Analysis of Microarrays (SAM) were performed using MeV software (27). HCL is a non-supervised analysis that allows grouping samples by similar expression patterns. All the identified proteins were used to build an HCL based on Pearson correlation. After HCL, a SAM was used to characterize differences between groups identified by HCL. On the other hand, SAM allows establishing differential expressed proteins between groups. This analysis consists in a t-test corrected by permutations over the number of samples. False Discovery Rate (FDR) was used to determine the significance (28).

### Probabilistic graphical models and functional node activities

The probabilistic graphical model (PGM) was built using *grapHD* package (29), R v3.2.5 and proteomics data without any a priori information. PGM are useful to analyze high-dimensional data. The analysis consists in two sequential steps: first, the searching of the spanning tree with the maximum likelihood, and, then, an edge depuration based on Bayesian Information Criterion (BIC) (30). The resulting network was divided in branches or functional nodes and, by gene ontology analyses, a main biological function for each branch was established. Gene ontology analyses were done using DAVID webtool (31) selecting “homo sapiens” as background and GOTERM-FAT, KEGG and Biocarta as categories. Functional node activity was calculated as the mean expression of these proteins that are related with the main function of each node, as previously described (32–34). Functional node activities were compared using Mann-Whitney tests.

### Predictor construction

BRB Array Tool was used to correlate protein expression with tumor relapse (35). These proteins were selected according to their p-values (p<0.01) and used to build a prognostic signature that classified patients into two groups (high- and low-risk) through a Cox regression. The predictor was internally validated using leave-one-out cross validation.

### Statistical analyses

Statistical analyses were done in GraphPad Prism v6 and SPSS IBM Statistics v20. P-values were considered as statistically significant under 0.05.

## Results

### Patient characteristics

One hundred and thirty-six patients with stage I to III TNBC were identified. One patient was excluded because no paraffin sample was available. Eight cases were excluded because they did not meet inclusion criteria (three with a follow-up under two years, four had received neoadjuvant therapy, and one had metastasis at diagnosis). Two samples were excluded due to the poor quality of the protein measurements. The final analyses included one hundred and twenty-five tumors.

Clinical characteristics are summarized in Table 1. This is an update of the clinical information included in our previous work (15). Median age was 56.8 years, 10% of the patients were younger than 40 years at diagnosis. Median follow-up was 64.6 months (1.1-257 months).

**Table 1:** Patients’ characteristics.

|  | Total (n=125) | Percentage (%) |
| --- | --- | --- |
| **Age at diagnosis. Median (range)** | 56.8 (24.7-85.2) |  |
| **Tumor size** |  |  |
| T1 | 44 | 35.2% |
| T2 | 67 | 53.6% |
| T3 | 7 | 5.6% |
| T4 | 7 | 5.6% |
| **Lymph node status** |  |  |
| N0 | 63 | 50.4% |
| N1 | 39 | 31.2% |
| N2 | 8 | 6.4% |
| N3 | 14 | 11.2% |
| Unknown | 1 | 0.8% |
| **Grade** |  |  |
| 1 | 3 | 2.4% |
| 2 | 17 | 13.6% |
| 3 | 103 | 82.4% |
| Unknown | 2 | 1.6% |
| **TNM stage** |  |  |
| I | 31 | 24.8% |
| IIA | 39 | 31.2% |
| IIB | 25 | 20.0% |
| IIIA | 10 | 8.0% |
| IIIB | 5 | 4.0% |
| IIIC | 14 | 11.2% |
| Unknown | 1 | 0.8% |
| **Treatment** |  |  |
| Anthracyclines and taxanes | 31 | 24.8% |
| Anthracyclines | 48 | 38.4% |
| Taxanes | 1 | 0.8% |
| Neither anthracyclines nor taxanes | 33 | 26.4% |
| None | 7 | 5.6% |
| Unknown | 5 | 4.0% |

Fifty-six (44.8%) patients had a relapse. Most of the relapses occurred in women with tumors greater than 2 cm, grade 3 and positive lymph nodes. Median follow-up of patients who did not relapse was 8.0 years (2.0-21.4 years).

Ten (18%) patients had a local and/or regional relapse, whereas 82% had a distant relapse (11 patients presented more than one location at first relapse) being the most frequent locations CNS and lung.

More than half of the relapses (n=33, 59%) occurred in the first two years since surgery of the primary tumor (median=20.2, range=1-135 months, median 22.8 months for local relapses and 18.6 for distant relapses) (Figure 1).

**Figure 1:**
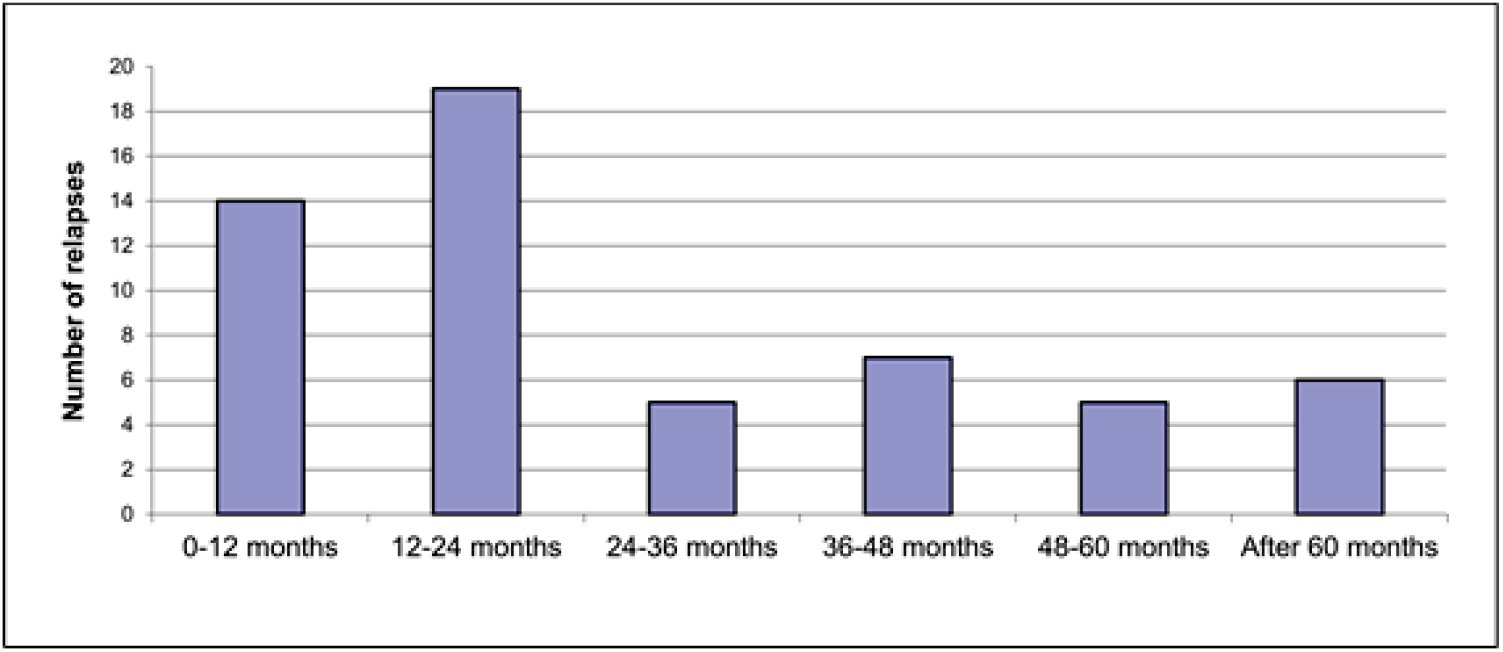
Time of relapse (months since surgery of the primary tumor to tumor relapse).

Seven patients did not receive any treatment after surgery (three of them, older than 80 years). Thirty-three patients received chemotherapy that did include neither anthracyclines nor taxanes (CMF was still widely used in the late 90s). Only six patients of those who received CMF had a relapse (11 patients with stage I, 21 with stage II, and only one patient with stage III).

### Proteomics experiments

As a first step towards the molecular characterization of TNBC tumors, we quantified the proteome of 140 formalin-fixed paraffin-embedded TNBC samples using the DIA+ acquisition method (20). This analysis enabled the quantification of 3,092 proteins, of which 1,206 proteins were consistently detected and quantified in at least 66% of the samples. Three samples were excluded from further analysis because they had significantly less quantified proteins (Figure 2). A gene ontology analysis showed that the identified proteins were mainly related with translational processes and extracellular membrane.

**Figure 2:**
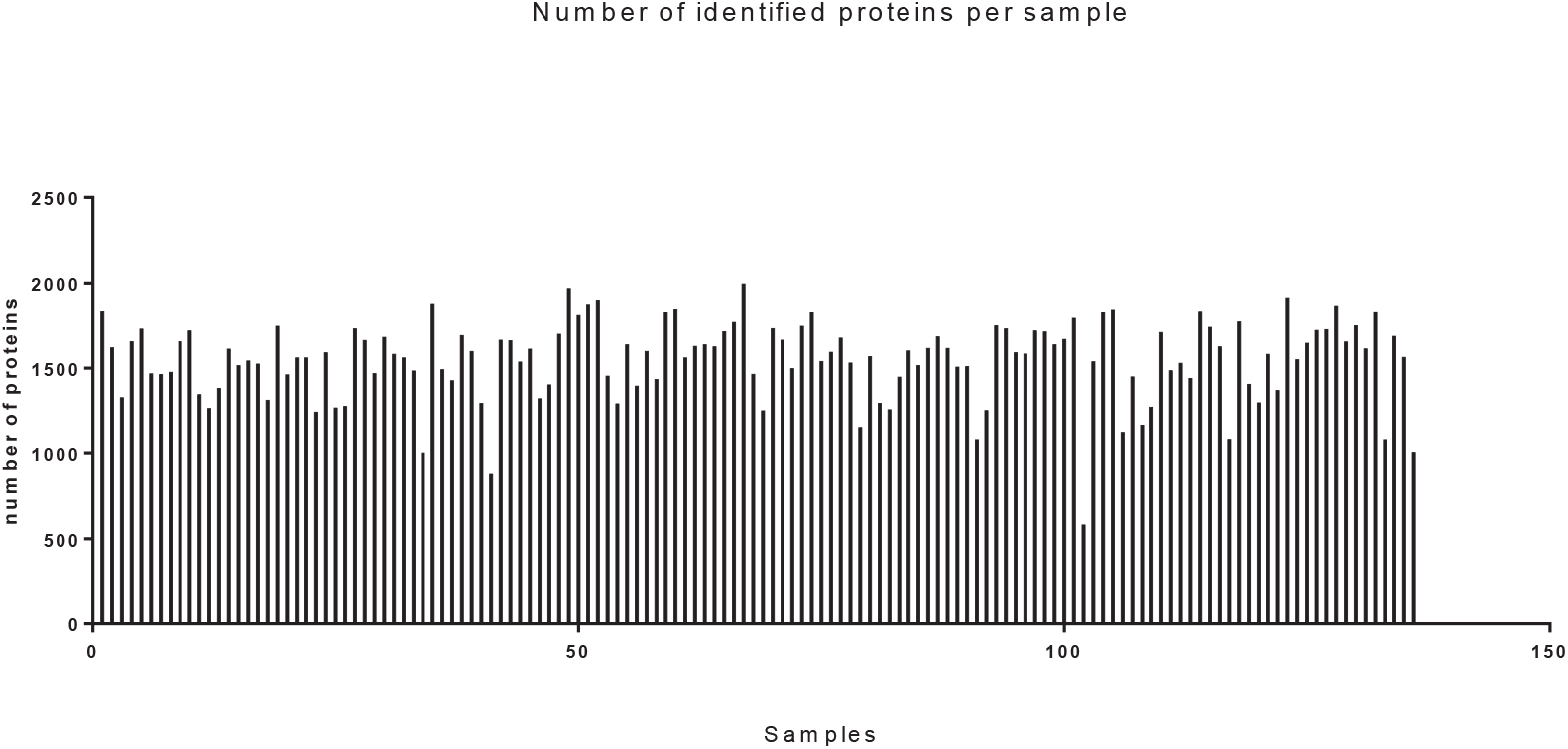
Number of proteins identified by sample using DIA+.

### Molecular classification

Using an unsupervised hierarchical cluster analysis (HCL), two different molecular groups were defined (Sup Fig 1). Group 1 included 76 patients and group 2 included 49 patients. However, we did not find significant differences in disease-free survival (DFS) between both groups (Sup Fig 2).

By SAM, 439 differentially expressed abundant proteins between the two groups were identified (Figure 3). Most of these proteins were related to membrane, adhesion, translation, glycolysis, and mitochondria. The main functions of the proteins more abundant in Group 2 were focal adhesion and membrane whereas those proteins predominant in Group 1 were related to mRNA translation and splicing, antigen presentation and T cells, focal adhesion and the anaphase process.

**Figure 3:**
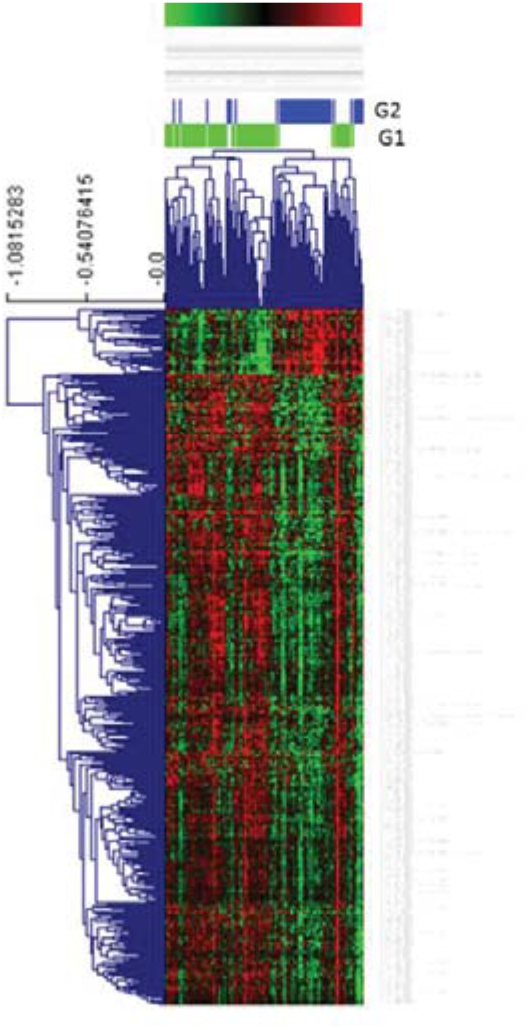
SAM between the two groups defined by HCL. G1= Group 1; G2= Group 2. Green= underexpressed, Red= overexpressed.

### Functional proteomics

A probabilistic graphical model (PGM) was built using the 1,206 quantified proteins to study protein functional relationships. The resulting network was divided into 10 functional nodes by gene ontology analyses: immune, cytoskeleton, glycolysis, transcription, mitochondria and oxidative phosphorylation, lysosome, splicing, exosome and two functional nodes related to cell adhesion (Figure 4).

**Figure 4:**
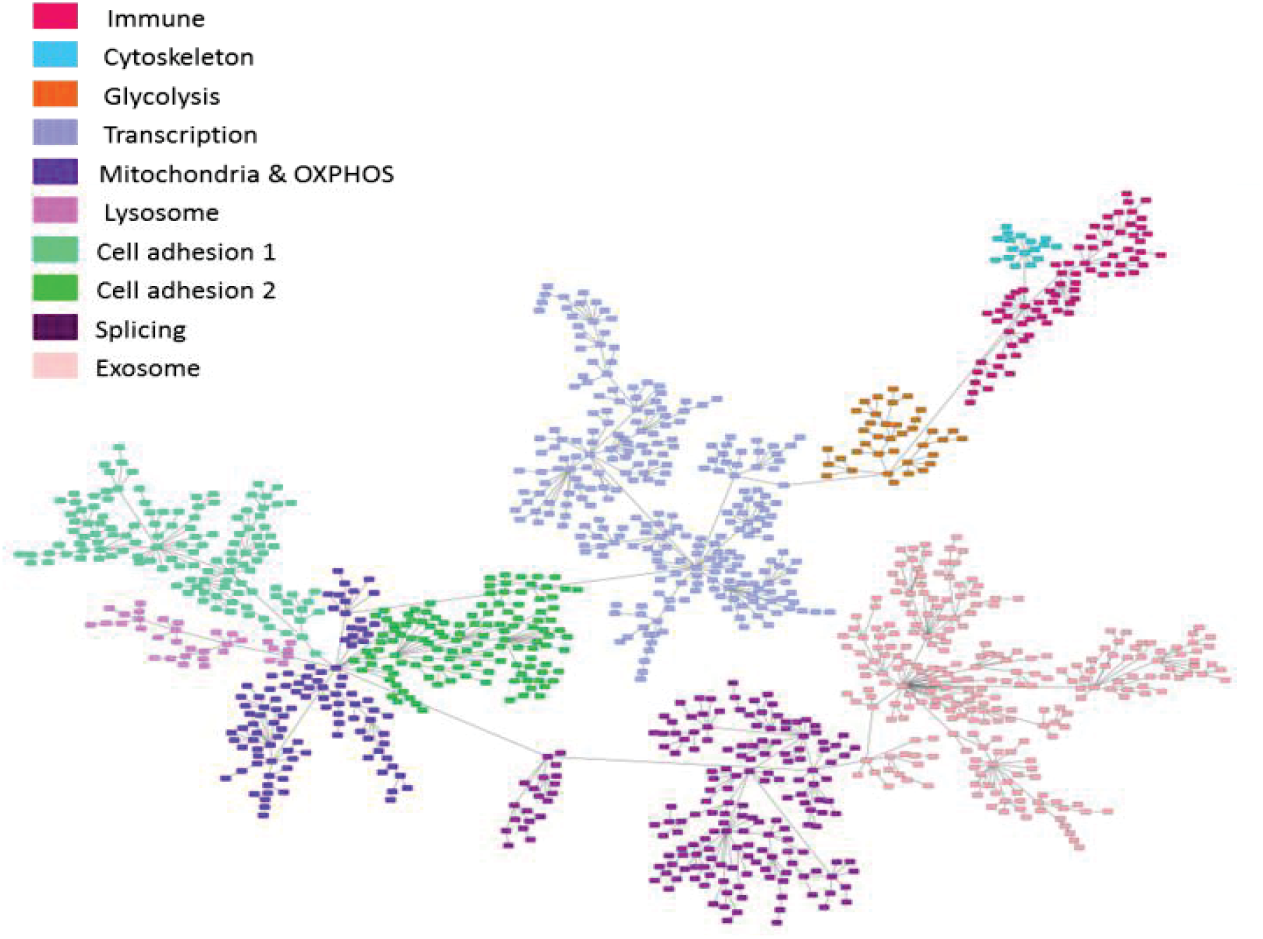
The probabilistic graphical model (PGM) functionally characterized built using the expression of the 1,206 identified proteins.

Functional node activities were then calculated as the mean of the expression of those proteins related to the biological function of each functional node. Comparing the two molecular groups, there were significant differences in glycolysis, immune response, extracellular matrix, exosomes, lysosomes, and cytoskeleton functional activities (Figure 5).

**Figure 5:**
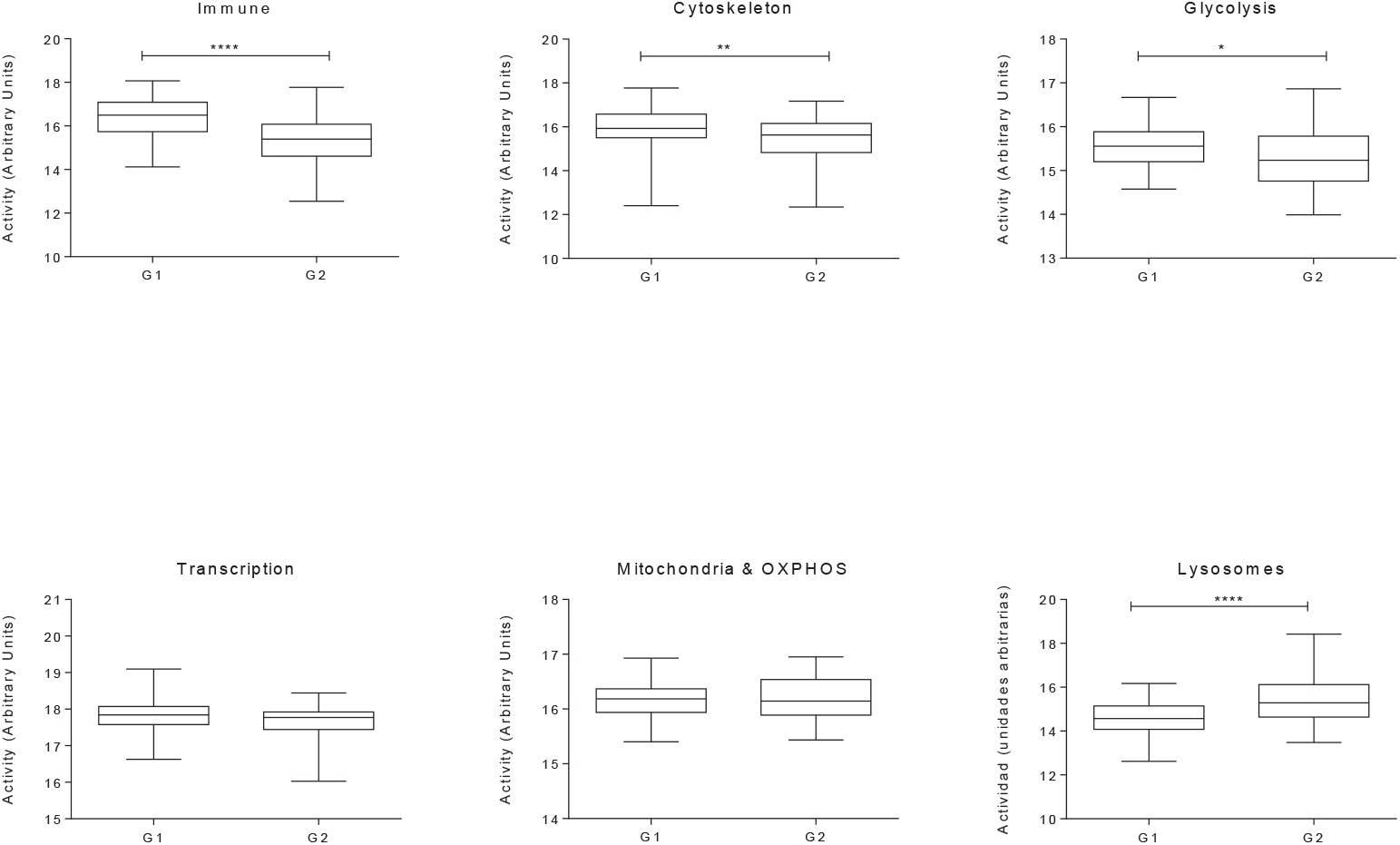
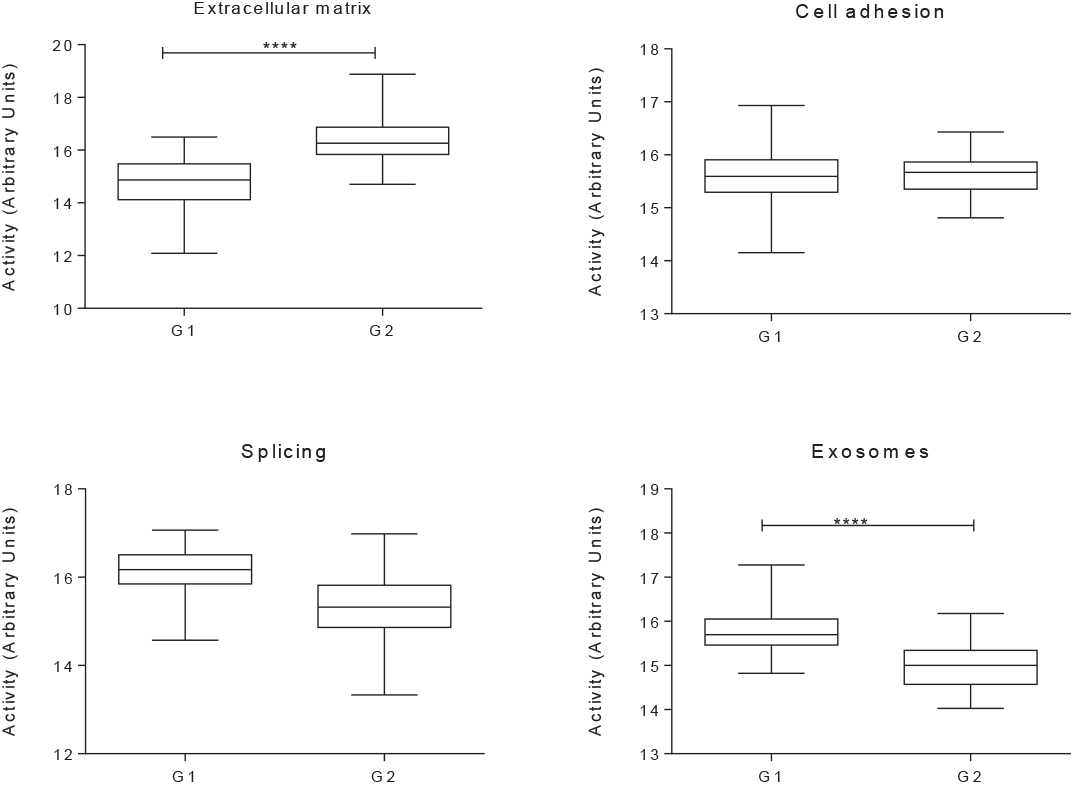
Functional node activities calculated for each molecular group. G1= group 1, G2= group 2. * p ≤ 0.05, ** p ≤ 0.01, *** p ≤ 0.001, **** p ≤ 0.0001.

### Proteins related with tumor relapse and disease-free survival signature

Next, proteins related to relapse and disease-free survival were identified and a prognostic signature was defined. Initially, twenty-nine out of 1,206 identified proteins were prioritized based on their association with relapse (p <0.01, Table 2).

**Table 2:**
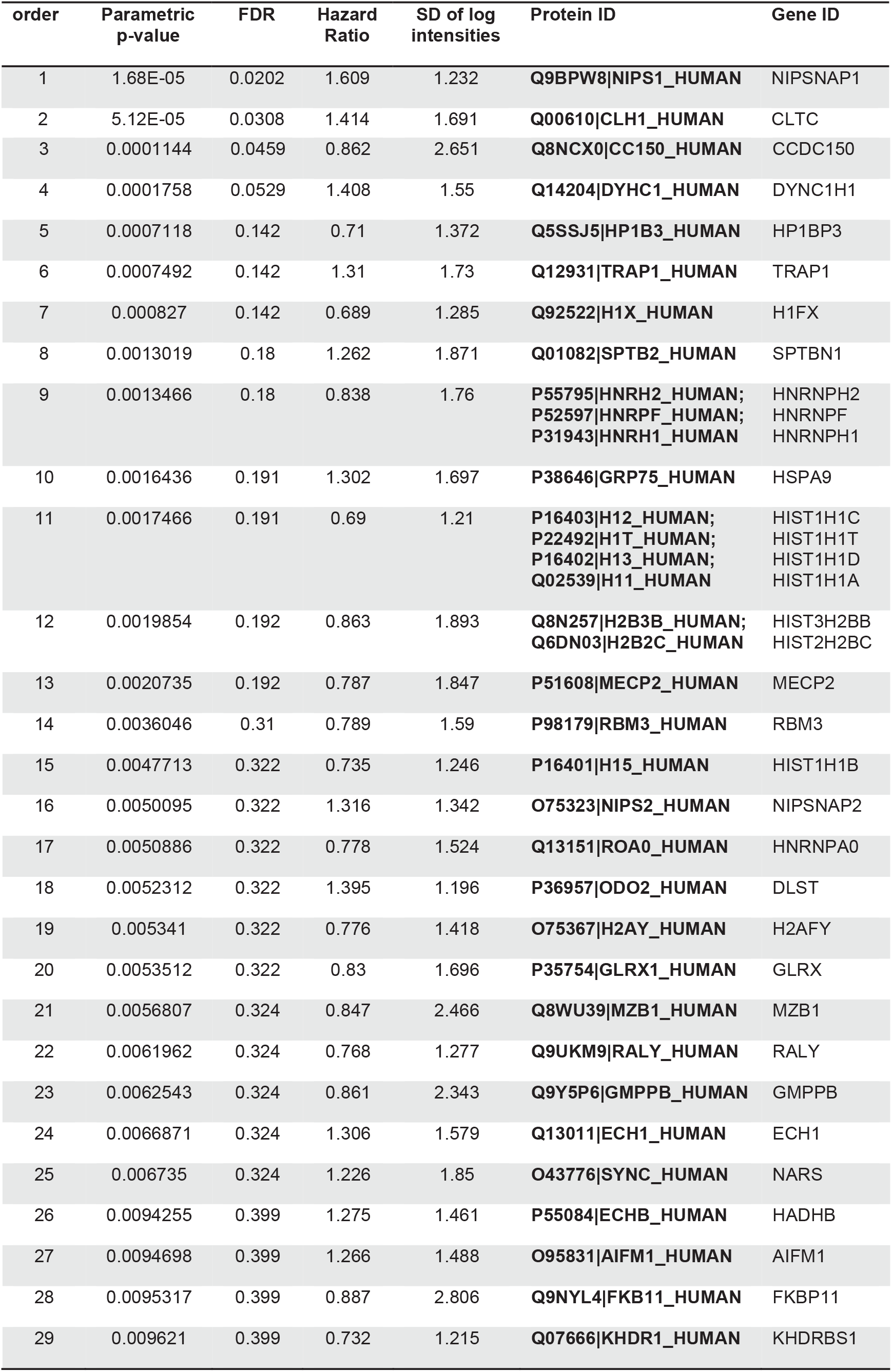
Proteins associated with relapse (p <0.01).

| order | Parametric p-value | FDR | Hazard Ratio | SD of log intensities | Protein ID | Gene ID |
| --- | --- | --- | --- | --- | --- | --- |
| 1 | 1.68E-05 | 0.0202 | 1.609 | 1.232 | Q9BPW8\|NIPS1_HUMAN | NIPSNAP1 |
| 2 | 5.12E-05 | 0.0308 | 1.414 | 1.691 | Q00610\|CLH1_HUMAN | CLTC |
| 3 | 0.0001144 | 0.0459 | 0.862 | 2.651 | **Q8NCX0\|CC150_HUMAN** | CCDC150 |
| 4 | 0.0001758 | 0.0529 | 1.408 | 1.55 | **Q14204\|DYHC1_HUMAN** | DYNC1H1 |
| 5 | 0.0007118 | 0.142 | 0.71 | 1.372 | **Q5SSJ5\|HP1B3_HUMAN** | HP1BP3 |
| 6 | 0.0007492 | 0.142 | 1.31 | 1.73 | **Q12931\|TRAP1_HUMAN** | TRAP1 |
| 7 | 0.000827 | 0.142 | 0.689 | 1.285 | **Q92522\|H1X_HUMAN** | H1FX |
| 8 | 0.0013019 | 0.18 | 1.262 | 1.871 | **Q01082\|SPTB2_HUMAN** | SPTBN1 |
| 9 | 0.0013466 | 0.18 | 0.838 | 1.76 | **P55795\|HNRH2_HUMAN;**<br>**P52597\|HNRPF_HUMAN;**<br>**P31943\|HNRH1_HUMAN** | HNRNPH2<br>HNRNPF<br>HNRNPH1 |
| 10 | 0.0016436 | 0.191 | 1.302 | 1.697 | **P38646\|GRP75_HUMAN** | HSPA9 |
| 11 | 0.0017466 | 0.191 | 0.69 | 1.21 | **P16403\|H12_HUMAN;**<br>**P22492\|H1T_HUMAN;**<br>**P16402\|H13_HUMAN;**<br>**Q02539\|H11_HUMAN** | HIST1H1C<br>HIST1H1T<br>HIST1H1D<br>HIST1H1A |
| 12 | 0.0019854 | 0.192 | 0.863 | 1.893 | **Q8N257\|H2B3B_HUMAN;**<br>**Q6DN03\|H2B2C_HUMAN** | HIST3H2BB<br>HIST2H2BC |
| 13 | 0.0020735 | 0.192 | 0.787 | 1.847 | **P51608\|MECP2_HUMAN** | MECP2 |
| 14 | 0.0036046 | 0.31 | 0.789 | 1.59 | **P98179\|RBM3_HUMAN** | RBM3 |
| 15 | 0.0047713 | 0.322 | 0.735 | 1.246 | **P16401\|H15_HUMAN** | HIST1H1B |
| 16 | 0.0050095 | 0.322 | 1.316 | 1.342 | **O75323\|NIPS2_HUMAN** | NIPSNAP2 |
| 17 | 0.0050886 | 0.322 | 0.778 | 1.524 | **Q13151\|ROA0_HUMAN** | HNRNPA0 |
| 18 | 0.0052312 | 0.322 | 1.395 | 1.196 | **P36957\|ODO2_HUMAN** | DLST |
| 19 | 0.005341 | 0.322 | 0.776 | 1.418 | **O75367\|H2AY_HUMAN** | H2AFY |
| 20 | 0.0053512 | 0.322 | 0.83 | 1.696 | **P35754\|GLRX1_HUMAN** | GLRX |
| 21 | 0.0056807 | 0.324 | 0.847 | 2.466 | **Q8WU39\|MZB1_HUMAN** | MZB1 |
| 22 | 0.0061962 | 0.324 | 0.768 | 1.277 | **Q9UKM9\|RALY_HUMAN** | RALY |
| 23 | 0.0062543 | 0.324 | 0.861 | 2.343 | **Q9Y5P6\|GMPPB_HUMAN** | GMPPB |
| 24 | 0.0066871 | 0.324 | 1.306 | 1.579 | **Q13011\|ECH1_HUMAN** | ECH1 |
| 25 | 0.006735 | 0.324 | 1.226 | 1.85 | **O43776\|SYNC_HUMAN** | NARS |
| 26 | 0.0094255 | 0.399 | 1.275 | 1.461 | **P55084\|ECHB_HUMAN** | HADHB |
| 27 | 0.0094698 | 0.399 | 1.266 | 1.488 | **O95831\|AIFM1_HUMAN** | AIFM1 |
| 28 | 0.0095317 | 0.399 | 0.887 | 2.806 | **Q9NYL4\|FKB11_HUMAN** | FKBP11 |
| 29 | 0.009621 | 0.399 | 0.732 | 1.215 | **Q07666\|KHDR1_HUMAN** | KHDRBS1 |

These 29 proteins associated with relapse were then used to build a prognostic signature. This predictor split the patient cohort into low- and high-risk groups (p-value= 0.0002, Hazard ratio [HR]=6.51, 20-80%). The predictor was based on the expression of proteins NIPSNAP1 (NipSnap homolog 1) and RBM3 (RNA-binding protein 3) (Figure 6). The formula to calculate the prognostic index is ∑ *w_i_ x_i_* − 4.264, where PI is the prognostic index, and x refers to protein abundances, w to the weights of each protein and i to the sample. A sample is being classified in the high-risk group when the prognostic index (PI) is higher than −0.584. Known characteristics of each protein are provided in Table 3.

**Figure 6:**
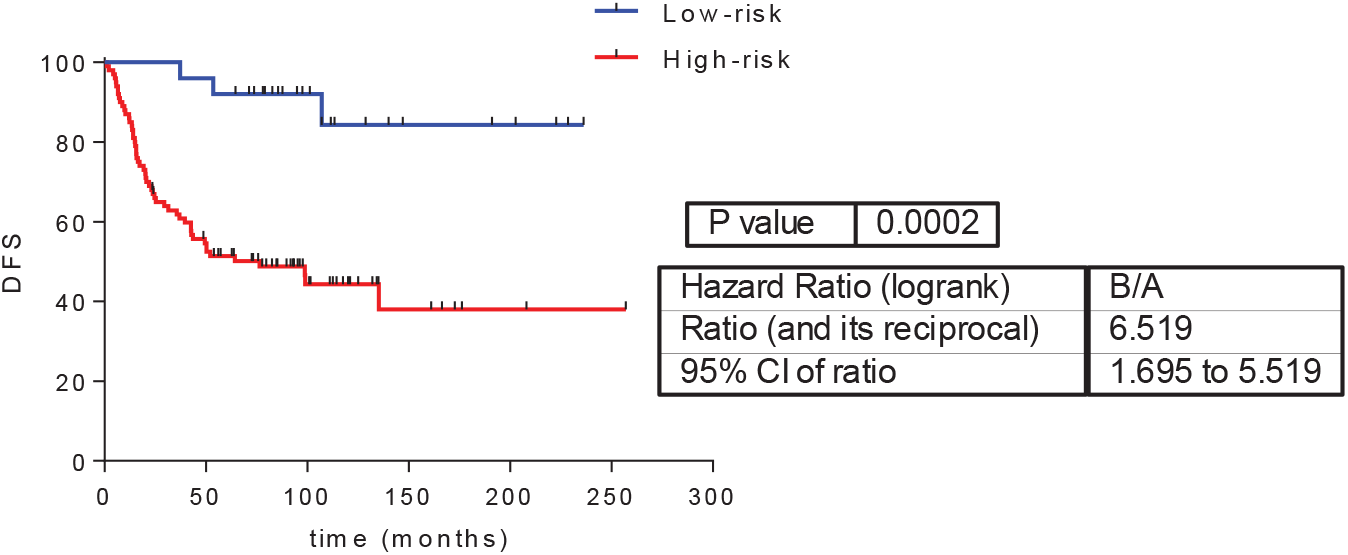
Predictor of relapse in TNBC based on the expression of NIPSNAP1 and RBM3. DFS = disease-free survival.

**Table 3:** Characteristics of the proteins identified by the predictor.

Characterization of triple negative breast cancer by DIA
| Gene ID | Protein name | Uniprot ID | Molecular function | Subcellular location |
| --- | --- | --- | --- | --- |
| NIPSNAP1 | Protein NipSnap homolog 1 | Q9BPW8\|NIPS1_HUMAN | - Neurotransmitter binding. | Mitochondria |
| RBM3 | RNA-binding protein 3 | P98179\|RBM3_HUMAN | - RNAm 3'-UTR binding.<br>- Poli(U) RNA binding.<br>- Ribosomal junction of large subunits<br>- RNA binding.<br>- RNAr binding of small ribosomal subunit.<br>- Translation repressor activity. | Nucleus |

Univariate and multivariate analysis of risk factors for tumor relapse are shown in Table 4. Tumor size, positive lymph nodes and the prognostic signature were significantly associated with an increased risk of relapse based on the univariate analysis. Multivariate statistical analysis showed that the prognostic signature was an independent prognostic factor for relapse.

**Table 4.** Risk factors associated with tumor relapse. HR= hazard ratio; CI= confidence Interval; P= p-value.

|  | Univariate Analysis |  | Multivariate Analysis |  |
| --- | --- | --- | --- | --- |
|  | HR (95% CI) | P | HR (95% CI) | P |
| *Grade* | 10.31 (0.01-10401.00) | 0.509 | --- | --- |
| *Tumor size (T)* | 1.92 (1.43-2.56) | <0.001 | 1.59 (1.14-2.22) | 0.007 |
| *Nodal status (N)* | 1.83 (1.45-2.30) | <0.001 | 1.47 (1.14-1.88) | 0.003 |
| *Prognostic signature* | 6.66 (2.07-21.37) | 0.001 | 5.19 (1.19-16.94) | 0.006 |

## Discussion

Breast cancer is the most frequent malignant tumor in women and among them, TNBC is associated with a worst prognosis. For this reason, most of patients with TNBC receive adjuvant or neoadjuvant chemotherapy, but it leads to overtreatment in some of them. Therefore, the identification of patients with low-risk of relapse would allow avoiding unnecessary treatments and toxicities. On the other hand, a better understanding of the underlying molecular biology could help to select those high-risk patients that could benefit from intensive follow-up and participation in clinical trials with new drugs. In this work, clinical and molecular features of a group of TNBC patients were analyzed.

Most of the patients included in this study presented grade III tumors, a size greater than 2cm, and lymph node involvement, similar characteristics to other TNBC patient series (1, 36, 37). Forty-five percent of the patients included in this study presented a relapse (local or distant). The local relapse rate was 8%, and the distant relapse rate 37%. In a similar study that included 83 TNBC patients treated with anthracyclines, distant relapse rate was 34% (36). In Dent et al., 34% of the patients experienced a distant relapse and 15%, a local relapse (1). TNBC has a greater incidence of visceral metastases than estrogen-receptor positive tumors (1, 6–9, 38), particularly at the CNS and lungs. In our cohort, CNS was the most frequent site of distant relapse (n=12, 10%), followed by lung metastases (n=10, 8%). Therefore, our sample was representative of the TNBC population.

Proteomics provides direct information about biological processes. Many efforts have been devoted to identifying proteomics signatures with prognostic value (15, 36, 39). Until now, proteomics based on mass-spectrometry has been the most used technique to identify therapeutic targets and prognostic biomarkers. However, despite the recent advances, only a few biomarkers have been identified and proteomics technology still has limitations for clinical application. The standard in proteomics experiments used to be LC-MS/MS using data-dependent acquisition (DDA) mode. Recently, data-independent acquisition methods have emerged, which enable the reproducible quantification and identification of proteins in large patient cohorts (40) with high accuracy and consistency(41).

A previous study using SWAT-MS in breast cancer samples established that TNBC was an heterogeneous group (22). In this study, fresh-frozen breast tissue samples were used to classify breast tumors in proteomics-based groups. A different study, which analyzed breast cancer cell lines of all subtypes and four TNBC tumor samples by LC-MS/MS, also established differences at protein level between breast cancer subtypes (42). Data obtained by our DIA experiments in breast cancer FFPE TNBC samples allowed us to divide patients with TNBC into two different groups (with 76 and 49 patients, respectively). Although no difference in survival appeared between these two groups, this molecular classification might provide patient stratification and be the basis for a targeted therapy for each one of these groups. The SAM found 439 proteins differentially expressed between these two molecular groups. Proteins were related to glycolysis, membrane, adhesion, mitochondria and translation. Molecular characterization of tumors is useful to define common biological alterations within subsets of patients that may become therapeutic targets. For example, SAM analysis showed that one of these differential processes is mitochondrial function. Metformin has an action on mitochondria and has been shown to affect cell viability in TNBC cell lines, so it could be useful in tumors that overexpress proteins related to mitochondria (43). Glycolysis was another relevant process, for which targeted drugs such as 2-D-deoxy-glucose are also available (44).

With the aim of studying protein relationships in TNBC, a PGM was built with the 1,206 protein abundances. This allowed us creating a graphical representation with ten functional nodes. There were differences in functional node activities between the two proteomics molecular groups in glycolysis, immune response, extracellular matrix, exosomes, lysosomes, and cytoskeleton. The characterization of differences at immune level has acquired a great relevance with the advent of immunotherapy. Differences in the immune functional node could be related to immunotherapy response. Recent studies suggests some role of anti-PD1 therapy in TNBC (45). Therefore, it would be interesting to find a good biomarker in this scenario to select patients for immunotherapy. Strikingly, there were significant differences in glycolysis functional node between the two groups that also appeared in SAM analyses. This functional approach offers complementary information to conventional analyses.

Moreover, in this work we have achieved the identification of a prognostic predictor based on proteomics data in TNBC. Twenty-nine proteins were related with relapse and the analysis of these proteins allowed us to build a prognostic signature based on two proteins: RBM3 and NIPSNAP1. RBM3 was associated with a lower risk of relapse, and NIPSNAP1, with worse prognosis. This protein signature is simpler and better, based on the obtained HR, than the P5 protein signature presented in previous works (15).

RBM3 is a member of the cold-shock protein family that regulates mRNA metabolism and has pleiotropic effects in cellular stress and oncogenesis (46). RBM3 protein is rarely overexpressed in normal tissues, but it is overexpressed in some solid tumors and, in recent studies, its expression levels seem to be related to prognosis and cytostatic sensitivity. High levels of RBM3 are an independent prognostic factor for DFS and overall survival in breast cancer (47, 48), ovarian carcinoma (49), gastric cancer(50), colon cancer (51, 52), prostate cancer (53), and melanoma (54). Our study confirms that RBM3 maintains its prognostic value in tumors with negative hormone receptors.

NIPSNAP1 is a protein usually expressed in CNS, liver and kidney. Its function is not clearly defined (55, 56). NIPSNAP1 is only expressed in neuronal tissues and it has been previously related to Alzheimer disease and phenylketonuria. Its association with cancer remains undefined. However, the fact that in our study, which comprehends a cohort with a significant presence of CNS metastases, NIPSNAP1 expression was associated with high-risk of relapse, suggests that it deserves future studies.

Despite the interest of our results, this study does not come without some limitations. First, it is worth mentioning that this was a retrospective study, and some patients received chemotherapy combinations that differ from the current standard. Moreover, the prognostic signature defined for predicting the risk of relapse is still a candidate prognostic signature and it will need prospective independent validation before its potential clinical application.

Overall, data-independent acquisition mass-spectrometry (DIA-MS) has demonstrated its utility in molecular characterization of archived triple negative breast cancer (TNBC FFPE) samples. In this study, we established two different molecular groups in TNBC patients with differential abundance of proteins related to mitochondria, membrane, adhesion and translation. The use of probabilistic graphical models (PGM) allowed the study of differences in biological processes between groups of patients and also suggested some processes with different activity between the two molecular groups. These processes could be exploited in the future as potential new therapeutic targets. In addition, proteomics data analysis allowed us to build a prognostic signature in TNBC population based on RBM3 and NIPSNAP1 abundances and the relationship between RBM3 and low-risk of relapse previously shown in other studies was confirmed.

## Acknowledgements

This work is supported by PI12/00444 from Instituto de Salud Carlos III, Spanish Economy and Competitiveness Ministry, Spain. The CRG/UPF Proteomics Unit is part of the Spanish Infrastructure for Omics Technologies (ICTS OmicsTech) and it is a member of the ProteoRed PRB3 consortium which is supported by grant PT17/0019 of the PE I+D+i 2013-2016 from the Instituto de Salud Carlos III (ISCIII) and ERDF. We acknowledge support from the Spanish Ministry of Science, Innovation and Universities, “Centro de Excelencia Severo Ochoa 2013-2017”, SEV-2012-0208, and “Secretaria d’Universitats i Recerca del Departament d’Economia i Coneixement de la Generalitat de Catalunya” (2017SGR595). This project has also received funding from the European Union’s Horizon 2020 research and innovation program under grant agreement No 823839 (EPIC-XS). LT-F is supported by the Spanish Economy and Competitiveness Ministry (DI-15-07614). GP-V and MIL-H is supported by the Consejería de Educación, Juventud y Deporte of Comunidad de Madrid (IND2017/BMD7783). AZ-M is supported by Consejería de Educación e Investigación de la Comunidad de Madrid (IND2018/BMD-9262). EL-C is supported by the Spanish Economy and Competitiveness Ministry (PTQ2018-009760). The sponsors were not involved in the study design, in data collection and analysis, in the decision to publish or in the preparation of this manuscript.

## Data availability

The raw proteomics data have been deposited to the PRIDE repository (57) with the dataset identifier PXD021491.

## Abbreviations

AGC: auto-gain control
BIC: Bayesian information criterion
CNS: central nervous system
DDA: data-dependent acquisition
DFS: disease-free survival
DIA: data-independent acquisition
FDR: False discovery rate
FFPE: formalin-fixed paraffin-embedded
HCD: higher-energy collisional dissociation
HCL: hierarchical cluster
PGM: Probabilistic graphical model
SAM: Significance analysis of microarrays
TNBC: triple negative breast cancer

